# Paper-based Sensing of Fucosylated Biological Compounds

**DOI:** 10.1101/2020.10.30.362889

**Authors:** Fatima Enam, Emily Kramer, Frederick Robinson, Andrea Alvarez-Acosta, Rebecca Cademartiri, Thomas J. Mansell

## Abstract

Advances in sensing technology have enabled rapid analysis of various biomolecules including complex carbohydrates. However, glycan analysis is limited by the throughput and complexity of assays for quantifying them. We describe a simple, low-cost enzymatic assay for the rapid analysis of fucosylation, down to linkage specificity, and its application to high-throughput screening of biologically relevant fucosylated compounds, to facilitate simple and straightforward analytical techniques. Paper-based devices integrate biosensor platforms and other diagnostic assays by fusing them with wax printing technology, making their fabrication even more inexpensive and simple. The specificity of the assay is established by linkage-specific glycosidic enzymes and the colorimetric output is visible to the naked eye, with costs that are lower than fluorescence/luminescence-based assays ($0.02/reaction). This platform was further improved by enhancing storage stability to retain analytical performance over time using desiccation and freeze-drying techniques. The assay platform allows analysis of hundreds of samples in minutes and we anticipate that this rapid and simple analytical method will be extended towards developing a universal glyco-barcoding platform for high throughput screening of glycosylation.

## Introduction

L-fucose is a common terminal sugar in glycans and oligosaccharides that has been attributed to several key metabolic and structural functions^1,2^. It is found as a terminal modification on N-, O- or lipid-linked oligosaccharides and as a core modification on N-glycans or linked to serine (Ser) or threonine (Thr) on proteins. It is also the second most abundant terminal monosaccharide moiety, after sialic acid^3^, found in mammalian carbohydrates to form various recognition motifs on many glycoproteins and glycolipids. Most notably, fucose has important roles as a component of ABO blood group antigens^4^, a key modulator of interactions in the immune system and in human breast milk.

Breastfeeding is associated with a lower risk of infection in babies and the underlying mechanism has been linked to the diverse range of complex sugars^5,6^, the human milk oligosaccharides (HMOs)^7^. A large majority of these HMOs are fucosylated^8,9^, with 2’-fucosyllactose (2’-FL) being the most abundant HMO comprising approximately 2 g/L. Notably, fucosylated HMOs inhibit binding and colonization of pathogens^10,11^ and provide protection against diarrhea associated with stable toxin of *Escherichia coli^12,13^*. Supplementation of 2’-FL to infant diet showed improved outcomes on innate and adaptive immune profiles^14^, and increasing evidence of its health benefits has led to its commercialization as a supplement in infant formula.

Similarly, the lipopolysaccharide (LPS) of *Helicobacter pylori* strains expresses Lewis antigens (Le^x^, Le^y^, H type I blood group structures) resembling the surface of host gastric epithelial cells^15^ and upon *H. pylori* infection, this may cause antibodies to bind not only to the bacteria but also the host epithelium resulting in tissue destruction. This antigen mimicry, being analogous to ABO blood group antigens, may also provide persistence through immune evasion^16^. Fucosylation is also an important oligosaccharide modification involved in cancer and inflammation^17^. The majority of human immunoglobulins (IgG) contain core fucosylation of the N-glycan in the Fc region, the absence of which has shown to have enhanced antibody-dependent cellular cytotoxicity (ADCC) activity due to increased binding affinity^18,19^.

Despite the biological importance of fucose, a critical obstacle in elucidating its biochemical roles lies in the technical difficulties associated with detection and analysis, especially with determining carbohydrate linkage positions. Current state-of-the-art methods for determining fucosylation include capillary electrophoresis (CE), mass spectrometry (MS), HPAEC-PAD^20^, NMR^21^ or glycan sequencing, which are laborious methods requiring specialized equipment and sample preparation. Another established method with higher throughput is the spectrophotometric detection of free L-fucose following defucosylation by measuring the formation of NADH^22^. Seydametova et al. demonstrated quantitative assays for 2’-FL, utilizing hydrolysis by fucosidases^23,24^. Their efforts, however, focused on a single biological molecule and by spectrophotometrically measuring the end-point nicotinamide adenine dinucleotide phosphate (NADP^+^) formed after an hour. Advances in synthetic biology have enabled high-throughput screening via biological components that have been standardized, for example, whole cell biosensors allow for the transduction of analyte concentrations to changes in gene expression that facilitate high-throughput assaying^25–27^. We previously demonstrated a genetically encoded whole cell biosensor to facilitate metabolic engineering by enabling the linkage-specific high-throughput detection of 2’-FL and four other HMOs^28^. The study demonstrated orthogonal screening of HMO structures, but entails culturing bacterial strains relatively expensive equipment to read fluorescence, e.g., flow cytometer or microplate reader.

To leverage the specificity and sensitivity of this assay to a faster and cheaper format, we sought to transfer the underlying detection mechanism to a paper-based assay. Paper based devices can be preferable to whole-cell assays, especially at the point of care, owing to their simplicity, low-cost and portability and the minimal resources, time and equipment required in their fabrication^29^. These devices also allow for rapid analysis of multiple samples in parallel. Colorimetric assays producing a visual readout are a common choice for paper-based devices^30–32^, with their usefulness lying in the formation of brightly colored formazan products upon reduction. Notably, they can be particularly useful in point-of-use or low-resource settings.

Here, we demonstrate a standard colorimetric assay for a simple, rapid, low-cost linkage-specific sensing system for fucosylated biomolecules (**Figure 1A**). To create this device, we expressed enzymes specific for three of the most common fucose linkages heterologously in *E. coli* and immobilized them on to the paper device. Using a formazan reduction assay coupled with fucose dehydrogenase, we then assay for the release of free L-fucose. The assay has a limit of detection of below 20 mg/L for 2’-FL and can detect fucose linked to several types of biomolecules, including HMOs, Lewis structures, and N-glycans. Finally, the assay can detect fucosylated compounds in complex body fluid matrices such as human milk and simulated urine.

**Figure 1:**
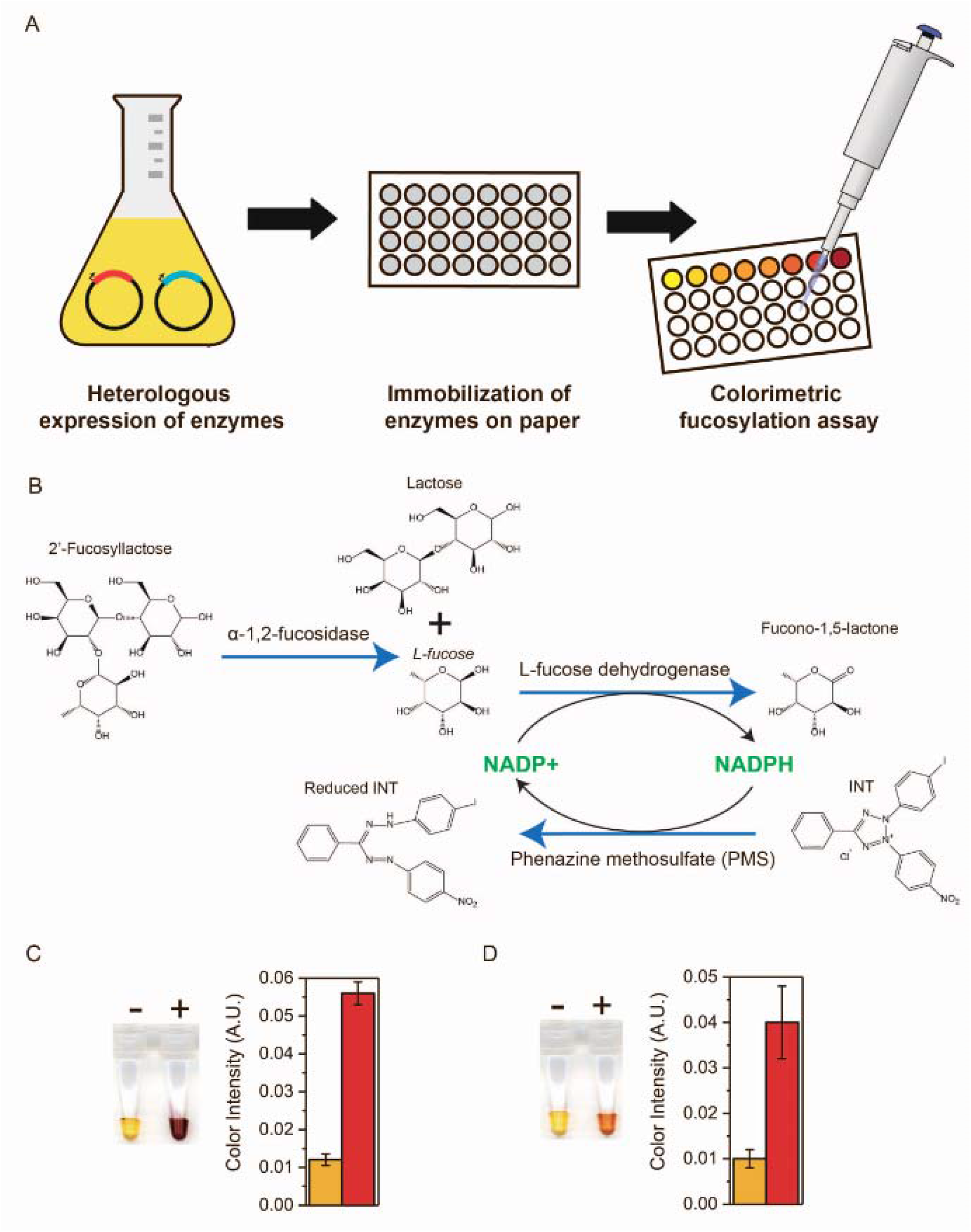
Strategy for paper-based sensing of fucosylation. **(A)** Schematic illustrating the immobilization of the enzymes on to the paper device and development of colorimetric signal upon addition of analyte. **(B)** Reaction mechanism for detection of fucose using a linkage specific fucosidase coupled to a colorimetric assay based on INT tetrazolium dye, 2’-FL shown here. **(C)** Images of reaction with free fucose (left) and 2’-FL (right) at 300 mg/L.

## Results

### Fucosidases act as linkage-specific detection elements for fucosylated HMOs

To couple reduction of colorimetric dye to the quantification of 2’-FL, we used fucosidases known in literature^33,34^ to be specific to the carbohydrate linkage to fucosylated molecules. Free fucose can then be assayed using a redox-coupled reaction in which fucose dehydrogenase (FDH) oxidizes L-fucose to fuconolactone, creating NADPH, which reduces the formazan dye. To confirm that this coupled reaction scheme would produce NADPH, we incubated 2’-FL (Fuc-α-1,2-Gal-β-1,4-Glc) with commercial purified α-1,2-fucosidase to cleave the fucose and in the presence of commercial FDH and NADP^+^. By following the increase in absorbance at 340 nm due to reduction of NADP^+^ on a microplate reader, we verified the redox reaction undergone by the cleaved fucose (**Supplementary Figure S1A**).

However, the use of two purified enzymes can be expensive. To achieve an efficient, low-cost detection system, we hypothesized that using a bacterial host as a heterologous system for the production of high levels of functionally active recombinant enzymes would be cost-effective. Because it efficiently and orthogonally cleaved L-fucose from 2’-FL in the context of a whole-cell biosensor^35^ we chose to express the soluble glycosyl hydrolase domain of AfcA, an α-1,2-fucosidase from *Bifidobacterium bifidum^36^* under a constitutive promoter, to liberate fucose from 2’-FL^35^. Cell lysate from overnight cultures of *E. coli* BL21 (DE3) expressing AfcA was added to purified FDH with NADP^+^ as before. We detected reduction to NADPH when 2’-FL was added as an analyte but not on the isomeric HMO 3-FL (Galβ-1,4(Fuc-α-3)Glc) (**Supplementary Figure S1A**). No background redox activity associated with the components in the cell lysate itself was observed in an empty vector control over a period of one hour. This strain served as a control in subsequent experiments, where the background noise from the crude cell lysate was noted for all measurements.

### Tetrazolium dye allows colorimetric detection of fucosylated compounds

Once we confirmed that the sequential reactions of cleavage by fucosidase followed by oxidation by fucose dehydrogenase generated the reducing co-factor, we added a redox-sensitive dye to make the assay visible by the naked eye. Tetrazolium salts have been historically used for testing cell viability, proliferation and cytotoxicity^37,38^. We chose 2-(4-iodophenyl)-3-(4-nitrophenyl)-5-phenyl-2H-tetrazolium chloride (INT), which in the presence of the intermediate electron carrier, 5-methyl-phenazine methyl sulfate (PMS), is reduced to an insoluble formazan. This precipitate absorbs light at a wavelength of 500 nm and provides a robust signal visible to the naked eye^39^. The generated NADPH, when coupled to INT, forms a stable red formazan (**Figure 1B**) and, being stoichiometrically proportional to the fucose concentration allows us to directly quantify the fucose. We determined the optimum pH and buffer conditions for the enzyme cocktail in the assay and coupled it with the INT/PMS assay to establish a colorimetric assay protocol.

We first sought to quantify free fucose with FDH from cell lysate. An intense color change was observed with 500 mg/L L-fucose resulting in a 100% increase in absorbance within a few seconds as read on a plate reader at 500 nm (**Figure 1C**). The reaction contains 2.5 μg of L-fucose with NADP^+^ present in excess. In contrast, by measuring the NADPH formation by reading the absorbance at 340 nm, a 30% increase in absorbance was seen at equivalent L-fucose concentrations, which is to be expected since the molar extinction coefficient of INT formazan is 18000 M^−1^cm^−1^ compared to 6220 M^−1^ cm^−1^ for NADPH. Thus, the colorimetric system showed greater sensitivity than changes in NADPH readout as demonstrated by the slopes in **Supplementary Figure S3A**. Notably, minimal background was again observed with the empty vector control. In addition, the switch to a colorimetric reaction allowed us to scale down the reaction volume to 20 μL.

To demonstrate the utility of the assay in reliably detecting fucosylated molecules, we used the assay to detect 2’-FL using additional crude cell lysate from cells expressing pAfcA. We titrated the amount of cell lysate needed for optimum signal. For each 20 μL reaction, 5 μL lysate was added. This also exhibited strong absorbance, a four-fold increase, with 2’-FL (300 mg/L) at the same concentration of L-fucose (300 mg/L), containing, as previously tested (**Figure 1D**). The linkage specificity was validated by the lack of significant increase in absorbance when tested with its isomer 3-FL, indicating that the α-1,3 linkage was not defucosylated. Similarly, we went on to demonstrate that this method can be extended to the detection of α-1,3 linked fucose as well by using lysate of cells harboring plasmid pAfcB expressing α-1,3-fucosidase under identical experimental conditions. We observed a 1.4-fold change in absorbance with 3-FL but no color change was observed with 2’-FL (**Supplementary Figure S1B**). Similar to our previous biosensors, this assay also demonstrated linkage specificity and the ability to distinguish between isomeric HMOs.

### Fabrication of paper-based platform

Once the colorimetric assay was developed, we focused on translating it to a paper-based assay. Paper-based devices are fabricated from cellulose because it has advantages of being low-cost, abundant, lightweight, biodegradable and biologically compatible^40,41^. In addition, its porosity and hydrophilicity offers wicking properties due to capillary action that can handle small volumes of liquid with no external pumping^42^. To create our device, we used Whatman Grade 3 chromatography paper and defined the hydrophilic area using wax-printing method owing to its ease of printing using commercially available printers. The design layout for the paper corresponded to the dimensions of a standard 96-well plate^43^. We chose this to accommodate distinct test zones for independent reactions that can be carried out in parallel, also allowing for the use of multi-channel pipettors for rapid assaying. Once printed, we melted the wax in an oven. To ensure fully formed barriers, we observed the bottom side of the paper. Prior to heating, no pattern is visible at the back. Upon heating, the wax melts and vertically penetrates through the porous fiber network^44^, which becomes visible on the bottom. This allows for the aqueous sample and reagents to be contained within spatially separated hydrophilic zones. There is also some lateral diffusion of the ink which reduces the diameter of the detection zone, but this was taken into account during the design to ensure appropriate dimensions. We designed the template with each well having a line thickness of 1 mm and internal diameter of 5 mm; after heating, the diameter was reduced to 4 mm. Once cooled, we further tested the hydrophobic barriers for leaks by addition of colored water and looking for any sort of bleeding outside of the well. If the liquid failed to contain, the paper was heated longer until no bleed-through was visible. The procedure is illustrated in **Figure 2** and detailed in the Methods. Once the device was ready to use, the enzymes were directly applied to the detection zones. A master mix containing NADP^+^ dissolved in Tris-HCl buffer (pH 9.5) was then added to wet the test zones, followed by the analyte to be tested. Immediately following that, the dye was applied and allowed to dry under ambient conditions before scanning and analysis (**Supplementary Figure S2**). In our experiments, it took less than 1 min from the addition of the dye for a detectable color change in the test zones to occur (**Supplementary Video S1**).

**Figure 2:**
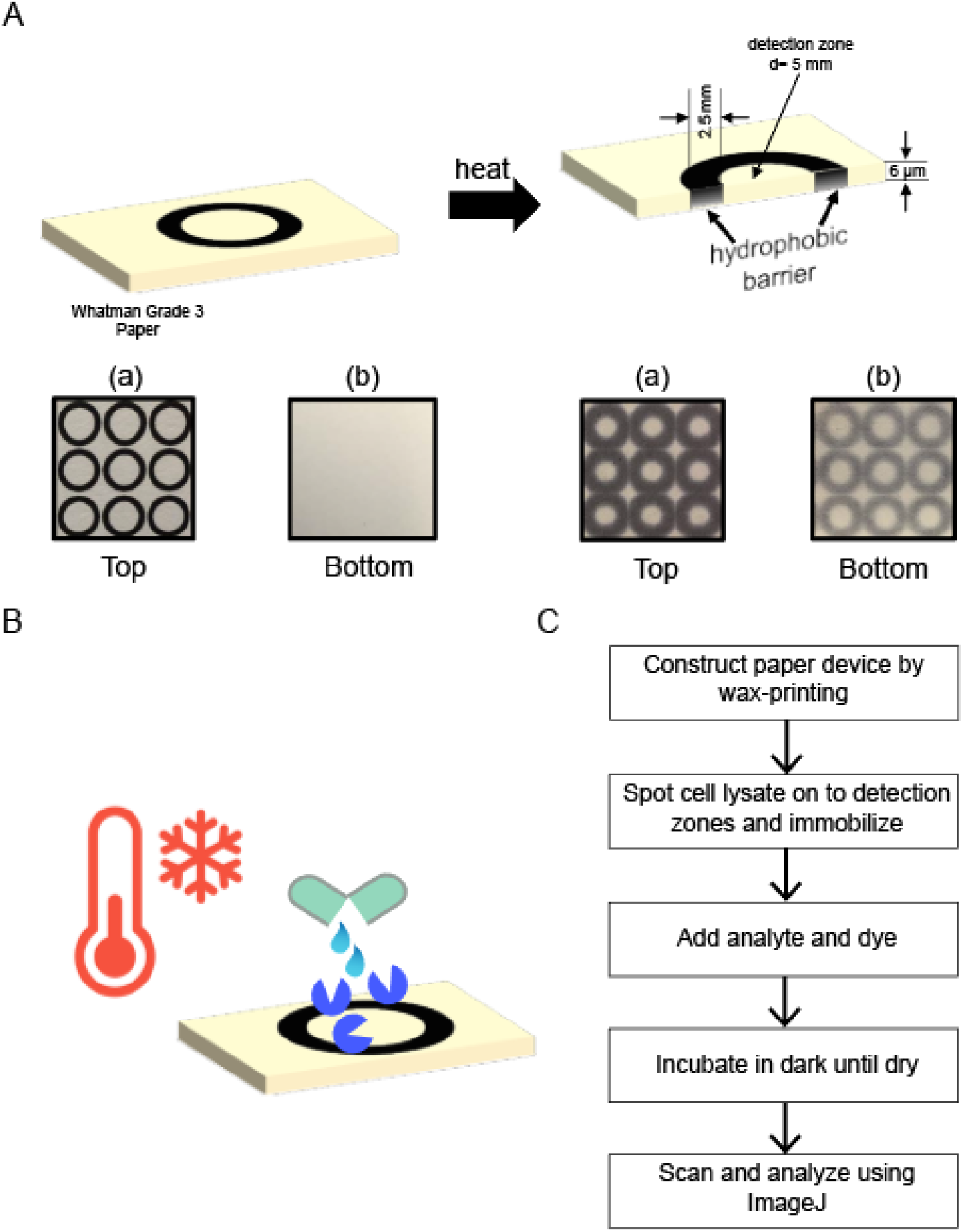
Schematic describing the design and process of setting up the paper-based assay. **(A)** Layout of the wax printed array. Top panel shows the dimensions of the wells. Bottom panel shows the printed array before and after heating, as seen on the top and bottom surface of the paper. **(B)** Cell lysate containing recombinantly expressed enzymes is spotted on the wells and immobilized. **(C)** Overview of the workflow of the assay.

### Paper-based sensing for distinguishing fucosylated HMOs

Next, we implemented this paper-based colorimetric assay for distinguishing three fucosylated HMOS: 2’-FL (Fuc-α-1,2-Gal-β-1,4-Glc), 3-FL (Galβ-1,4(Fuc-α-3)Glc) and DFL (Fuc-α-1,2-Galβ-1,4(Fuc-α-3)Glc). We also tested for GDP-L-fucose, the donor substrate, a key component in the biosynthesis of fucosylated HMOs^45,46^. Either synthetic routes for 2’-FL and 3-FL, enzymatic^47^ or microbial via metabolic engineering^45^, take place via GDP-L-fucose which can be present in the final product at different levels. It is imperative that our assay is not sensitive to GDP-L-fucose for its use in analysis of carbohydrate production from microbial sources. For the assay we spotted the required enzymes on to the paper: either no enzyme as control and for background reduction, only FDH for L-fucose and GDP-fucose detection, FDH and AfcA for 2’-FL detection, FDH and AfcB for 3-FL detection and combination of FDH, AfcA and AfcB for DFL detection. This was followed by the NADP^+^ dissolved in the buffer, the respective analytes and the dye. The paper was allowed to dry and for the color to fully develop for an hour at room temperature before analysis. We first tested if the two isomers, 2’-FL and 3-FL were distinguishable and if there was any crosstalk. We had established earlier the lack of substrate promiscuity in AfcA and AfcB in the context of the whole cell biosensor^28^. No fucose was liberated from 3-FL when AfcA was present (**Figure 3A**). **Figure 3B** shows a 25-reaction array to distinguish three fucosylated HMOs. The control zones (top row with no added enzymes and left column with no added substrate) showed no color change and the values obtained were noted to account for the background signal from the dye and improve the accuracy of the assay. Notably, no residual activity was observed with the concentrations of cell lysate used. An intense color change from yellow to red was observed with free L-fucose when FDH was present, as expected. This served as a positive control, confirming the activity of the FDH. With 2’-FL, a color change was observed only when AfcA was present to cleave the α-1,2-linked fucose. We thus obtained a signal only when AfcA or both AfcA and AfcB were present. We obtained similar results with 3-FL when AfcB was present. We did not observe any color change with GDP-fucose implying the lack of enzymatic activity of FDH on GDP-fucose as a substrate (data not shown).

**Figure 3:**
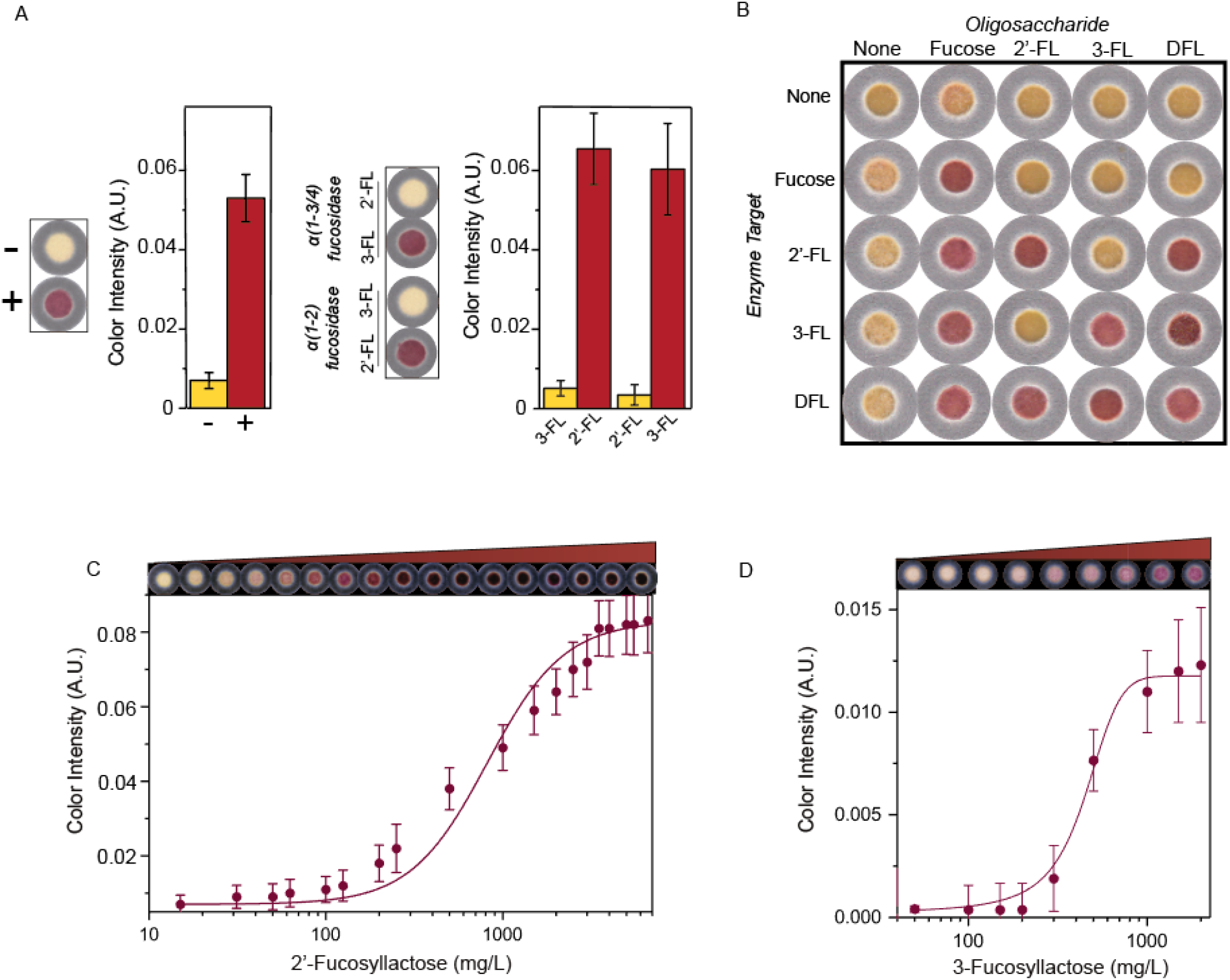
Determination of fucosylation in HMOs using paper device. **(A)** Reaction with free fucose (left) and fucosylated HMOs, 2’FL and 3-FL (right). Assays were performed with 500 mg/L of L-fucose, 2’-FL and 3-FL **(B)** A 25-reaction array with two different linkage specific fucosidases in different combinations. Image was captured after the paper was dry. **(C)** Transfer function for sensing 2’-FL. Response curve depicts mean color intensity (analyzed in triplicate). The dose-response curve was fitted with a 4PL model, shown by the line. The top shows a capture of the signal expression from increasing concentrations of 2’-FL. **(D)** Calibration curve for 3-FL, as described in (C). Error bars represent standard deviation of triplicate measurements of mean color intensities.

We generated the respective calibration curves by the addition of a series of dilutions of 2’-FL and 3-FL in range 0-6000 mg/L. We then scanned the paper device using a desktop scanner and analyzed the digital image on the green intensity channel. We quantified the color change in the test zones, and converted the color change value into the corresponding concentrations using the calibration curve for the assay (**Figure 3C, 3D**). The experimental data were fitted into a 4-parameter logistic model (4PL) using MATLAB and the resulting transfer functions revealed the dynamic range and limits of detection. The calibration curve for 2’-FL showed a wide dynamic range between 150 and 2500 mg/L of 2’-FL (0.3-5 mM) and the limit of detection (LoD) was found to be below 20 mg/L (0.04 mM) which satisfies the levels at which these biological molecules are produced either naturally or via synthesis. For example, Baumgärtner *et al.* engineered an *E. coli* strain to produce 2’-FL at levels of up to 21 g/L^45^ and analysis of HMOs in human milk shows 2’-FL to be present 60-15000 mg/L^48,49^, which is well within the limit of our device. Although the limit of detection is five-fold higher than our whole-cell biosensor, the dynamic range for this device is much wider. Similarly, we characterized the assay for 3-FL (**Figure 3D**) and the dynamic range of detection was found to be between 250 and 900 mg/L and the limit of detection below 250 mg/L (0.5 mM).

We also measured the absorbance values of the series of 2’-FL concentrations in a 96-well paper plate on a microplate reader to see the correlation with the values obtained from the paper-based assay. **Supplementary Figure S3B** shows the direct correlation of the absorbance values as measured with a plate reader with the values from the paper-based assay which correlate quite well.

### Stability of paper-based sensor

Since the intrinsic instability of enzymes or proteins in general, makes it challenging for them to be used for downstream applications, it was important to establish that our assay was functional over a longer period of time. Different strategies have been developed to produce more stable proteins and improve storage stability and shelf-life. The most frequently used method, especially in the biopharmaceutical industry is drying^50,51^. We spotted the enzymes on the test zones and either dried it at room temperature or by lyophilization. We then evaluated the stability of the enzymes on the cellulose matrix by storage at room temperature or at 4 °C. We performed the assay for 2’-FL detection under the four different conditions over a period of 45 days and imaged the wells to confirm expression of the enzymes (**Supplementary Figure S3C**). The desiccated enzymes maintained the similar activity over a period of one week, and slowly started to decline. In 5 weeks, the activity dropped to 50% of the original, under both storage conditions, and to 35% in 45 days. However, when lyophilized, 70% of its activity was retained after 45 days. It was noticed that with refrigeration, the decline in activity was slower compared to storage at room temperature. In solution, the enzyme master mix loses its activity at room temperature in one day and in less than a week at 4 °C. Overall, the data demonstrates sufficient stability of the paper-based sensor, especially under lyophilization. The assay can be stored at room temperature for shorter durations, avoiding the need for cold-chain. Beyond portability, immobilization of the enzymes enables minimal sample handling and cost and easy operation, without the need for specialized equipment and skills.

### Analysis of fucosylated HMOs in breast milk and maternal/infant urine

There is good evidence showing positive roles played by HMOs in breast milk, especially fucosylated HMOs, on infant health. The α-1,2 fucosyltransferase (FUT2) gene catalyzes the transfer of α-1,2-linked fucose residue, while the α-1,3/4 fucosyltransferase (FUT3) transfers α-1,3-linked fucose residues to HMOs. Secretor status (harboring a functional copy of the FUT2 gene) can be established by profiling of HMOs in breast milk^52^. This variation in the composition of HMOs in different mothers can provide some infants advantages over others. We tested if our assay was capable of detecting and quantifying 2’-FL and 3-FL in human breast milk to help identify secretors. As a control, we used powdered cow’s milk, reconstituted to mimic breast milk. We also chose to validate the quantitation using HPLC/MS-MS based on standard retention times and mass spectrometric analysis. For the human breast milk samples, we observed a color change indicating a concentration of 5,012 mg/L of 2’-FL (**Figure 4A**), which we confirmed with a series of dilutions, and consistent with the 5-15 g/L of HMO typically observed^49^. The signal from the paper-based assay indicating higher 2’-FL concentrations may be attributed to the presence of other α-1,2-fucosylated HMOs, such as lacto-N-fucopentaoses (LNFP I, II, III and V). The concentration of 3-FL was below the detection limit of the paper-based assay. However, we observed a color change once the samples were concentrated five and ten-fold (**Figure 4B**). 3-FL was found at 138 mg/L which corresponds to values found in literature^49^. We observed no color change with the cow’s milk samples, validating the robustness of the assay in measuring fucosylation in biological fluids. To validate the assay, characterization of isomers, 2’-FL and 3-FL, were carried out using mass spectrometry. Although we were able to get adequate separation between the 2’-FL and 3-FL peaks, there was overlap. The peaks were fully resolved by utilizing differences in m/z ratios of the deprotonated negative ions (MS) in conjunction with the difference in retention time (HPLC) to resolve the isomers (**Supplementary Figure S4**). To confirm the identity of the peaks, we also carried out an exoglycosidase (α-1,2-fucosidase or α-1,3-fucosidase) digest. Treatment with either α-1,2-fucosidase or α-1,3-fucosidase resulted in removal or major reduction of the expected peaks (**Supplementary Figure S5**). HMOs have also been detected in the serum and urine of mothers, with 2’-FL at the highest concentrations in secretors (20 mg/L)^53^. Studies have also shown high resemblance of HMO profiles in infant urine to mother’s breast milk, as some are absorbed and reach the systemic circulation and excreted in urine^54^. These samples can serve as excellent indicators of availability of fucosylated HMOs to infants, allowing for supplementation when needed. We spiked non-biological urine containing constituents that mimic human urine with 2’-FL within a physiologically relevant concentration range. The levels of 2’-FL in urine have been shown to correspond to levels in the milk. Goehring et al. found the urine of breast-fed babies of secretor mothers to contain 2’-FL in the range of 35-191 mg/L^55^. The assay responded to both low (20 mg/L) and high (200 mg/L) levels of 2’-FL spiked in Surine (**Figure 4C**). Importantly, we did not see any interference from the urine constituents with the enzymes in our assay.

**Figure 4:**
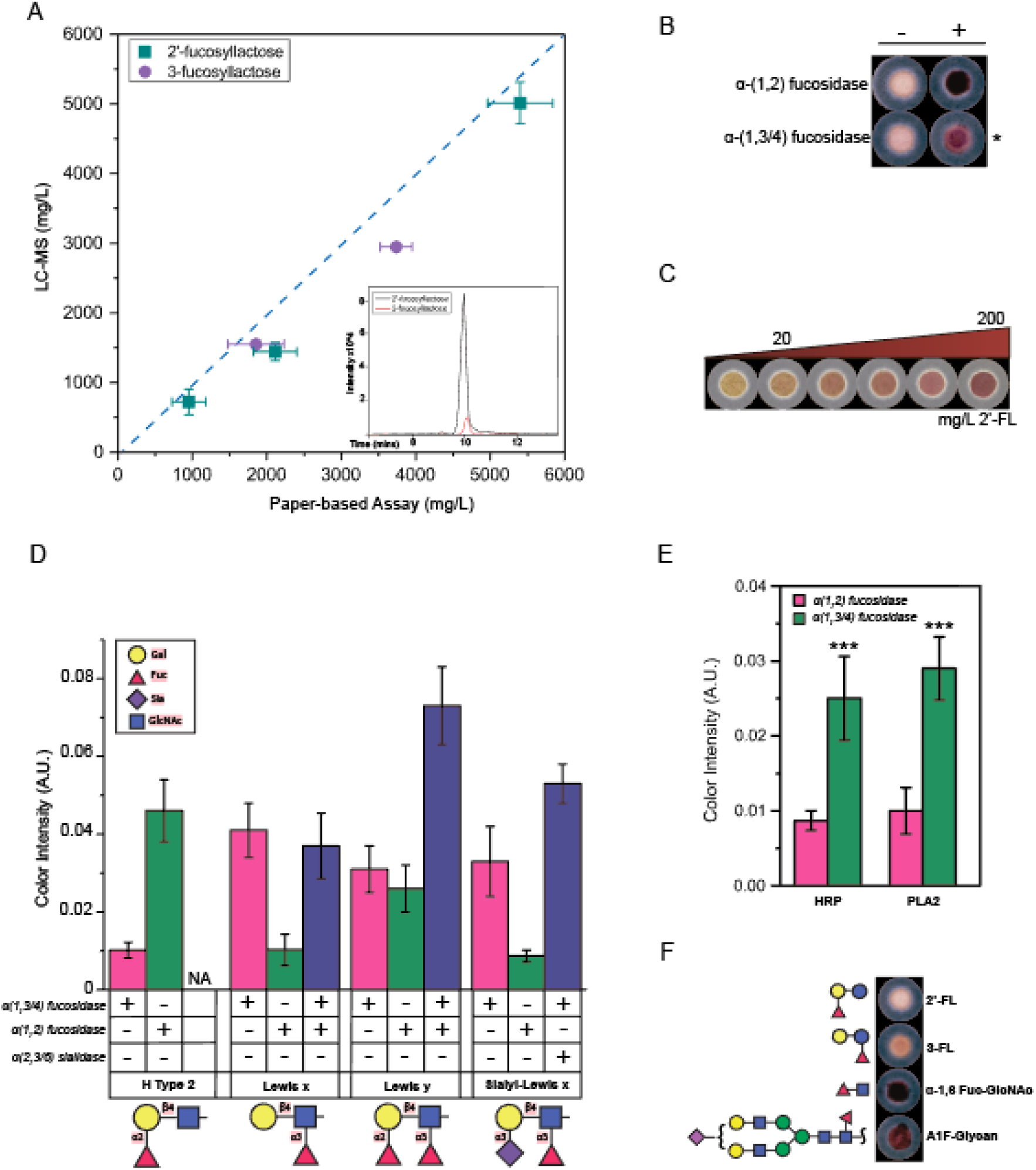
Paper-based assay enables determination of fucosylation type in biological fluids and glycoconjugates. **(A)** Comparing measurement of concentrations of 2’-FL and 3-FL in human breast milk using paper-based assay and HPLC/MS. Lipids and proteins were removed before analysis. To analyze 3-FL, samples were concentrated. Inset shows chromatogram peaks for 2’-FL (black) and 3-FL (red). **(B)** Arrayed image showing colorimetric output from assay on breast milk samples. * indicates concentrated samples tested. **(C)** Testing for 2’-FL in spiked synthetic urine at levels found in maternal and infant urine. **(D)** Distinguishing Lewis Type 2 antigens and quantification of their fucosylation. **(E)** α-1,3-Fucosylation in glycoproteins. Differences in the means of the test conditions and the controls were analyzed using a t-test (***P < 0.0001). **(F)** Determination of α-1,6-fucosylation in N-glycans. 2’-FL and 3-FL were used as controls.

### Determination of fucosylation in glycoconjugates

Glycoconjugates comprise a complex and diverse group of molecules that play important biological roles. Current methods of analysis of these molecules also involve labor-intensive procedures. We hypothesized that our paper-based scheme would be sufficient to analyze other fucosylated glycoconjugates. We targeted the highly fucosylated blood group Lewis antigens, specifically the Type 2 structures: composed of Gal-β(1,4)-GlcNAc, giving rise to Lewis^x^ (Le^x^), sialyl-Lewis^x^ (Sia-Le^x^) and Lewisy (Le^y^). We also tested the blood group H antigen, which is the fucosyl residue in blood group antigens. Due to the presence of either one or both of α-1,2/3/4-linked fucose, we took a combinatorial approach. With antigen H type 2, a colorimetric change was only observed with AfcA (**Figure 4D**). Similarly, Le^x^ was only detected in the presence of AfcB. With Le^y^ antigen, a three-fold increase in signal in the presence of two fucosidases was observed. Sia-Le^x^ is sialylated and we hypothesized that the sialic acid may interfere with the fucosidase activity. To test the hypothesis, we also introduced an α-2,3-neuraminidase that can cleave the sialic acid bound to the LacNAc. We observed a two-fold increase in the signal upon removal of sialic acid. Although the assay is not orthogonal, it allows us to easily distinguish the Lewis Type 2 antigens. This tool has the potential to determine the composition of HMOs in the milk of mothers as determined by the Lewis blood group and hence, further aid in the development of more personalized HMO supplementation. We next sought to determine if our paper-based assay had the capacity to detect fucosylation in the context of more complex biomolecules like glycoproteins. We targeted two common glycoproteins that are α-1,3-fucosylated: horseradish peroxidase (HRP) and phospholipase A2 (PLA2) to serve as models. Core α-1,3-Fuc is a common N-glycan modification found frequently on many plant and insect glycoproteins, but not mammalian ones. Since this core antigen does not occur in mammals it may be responsible for inducing immunogenic responses in mammals^56^. These experiments were consistent with our previous results; we were able to see a strong signal in the presence of α-1,3 fucosidase with both glycoproteins (**Figure 4E**).

One common mammalian N-glycan modification is core α-1,6 fucosylation. To specifically cleave α-1,6 fucosylated N-glycans, we produced AlfA from *Lactobacillus casei* by recombinant expression in *E. coli* as described earlier. AlfA is an α-1,6 fucosidase, shown to have specific hydrolytic activity on α-1,6 linked fucose^57^. The recombinant enzyme was tested and validated on 6'-fucosyl-GlcNAc and A1F-Glycan. 6’-Fuc-GlcNAc is an important modification of the core glycan in N-glycoproteins. A1F-Glycan is a monosialo-fucosylated biantennary oligosaccharide, found on different mammalian glycoproteins including IgG, gamma globulins, and many serum glycoproteins. We observed a strong color change with both substrates and corresponded well with the amount of fucose in the molecules (**Figure 4F**). The figures correspond to 0.4 μg and 2 μg of L-fucose in A1F-Glycan and 6’-Fuc-GlcNAc respectively. We also tested for the specificity of AlfA: no signal was seen with α-1,2/3 linked fucosylated substrates (2’-FL and 3-FL respectively). This indicates that the paper assay allows for efficient, linkage specific quantification of fucosylation in N-glycans. We attempted to detect α-1,6 fucosylation in IgG from human serum but did not see any signal, possibly due to fucose being below the limit of detection. No color change was observed when we tried analyzing the isolated N-glycans by treatment of IgG with endoglycosidase PNGase A/F.

## Discussion

As the role of carbohydrate structures in myriad biological processes is increasingly recognized, rapid and low-overhead analysis of carbohydrates becomes increasingly important. For example, fucosylation in serum alpha-fetoprotein can serve as a marker for early diagnosis of hepatocellular carcinoma^58^. However, current methods of characterizing glycans, especially to the level of linkage specificity, require expensive equipment and significant expertise. To enable more practical methods of diagnosis or development of glycoprotein therapeutics, advances in analytical glycomics are needed to overcome the bottlenecks associated with state-of-the art technologies. Here we demonstrate a robust, inexpensive, high-throughput colorimetric assay to quantify fucose in biologically relevant fucosylated molecules. The main advantage of this assay is easy and quick way to perform it, enabling in-line detection of multiple samples in parallel. This assay also minimizes the volume of sample needed for analysis. While this assay can be useful in low-resource settings, it also has the potential to leverage the metabolic engineering efforts for production of fucosylated glycans by enabling the high-throughput screening of libraries of mutant variants. This can also provide a new approach for directed-evolution efforts towards improved fucosyltranferase variants. In addition, integration of this method with other glycosyl hydrolases beyond fucosidase could be leveraged to provide comprehensive quantification of sugars and linkages in many complex carbohydrates. This platform demonstrates high specificity, reproducibility and stability, which constitutes a promising approach to be implemented as an easy-to-use protocol for sensitive and high-throughput detection of fucose in biological samples.

## Acknowledgements

This work was supported by Iowa State University Startup Funds. F.E. was funded by the NSF Trinect Fellowship and Manley Hoppe Professorship. T.J.M. was partially supported by the Karen and Denny Vaughn Faculty Fellowship. Frederick Robinson was supported by the NSF BioMaP REU program. The authors thank Dr. Ludovico Cademartiri for assistance with wax-printing. The authors also thank Dr. Lucas J. Showman at the W.M. Keck Metabolomics Research Laboratory for helping to analyze the human breast milk samples.

## Online Methods

### Chemicals and reagents

All reagents used were of analytical grade. Tris-HCl buffer (pH 9.5) was used as the buffer solution. 2-(p-iodophenyl)-3-(p-nitrophenyl)-5-phenyltetrazolium chloride (INT), phenazine methosulfate (PMS), NADP^+^ and dimethyl sulfoxide (DMSO) were purchased from Sigma-Aldrich (Millipore Sigma, St. Louis, MO). Luria-Bertani (LB) culture medium, kanamycin and carbenicillin were obtained from Sigma-Aldrich. Isopropyl b-D-1-thiogalactopyranoside (IPTG) was purchased from Invitrogen (Invitrogen, Carlsbad, CA). 2’-Fucosyllactose and 3-fucosyllactose were a kind donation by Glycom (Glycom A/S, Denmark). L-fucose, GDP-fucose, A1F-Glycan and 2-Acetamido-2-deoxy-6-O-(6-deoxy-α-L-galactopyranosyl)-D-glucopyranose (Fuc-α-1,6-GlcNAc) were purchased from Carbosynth (Carbosynth, Berkshire, UK).

Phospholipase A2 (*Apis mellifera*) was purchased from Enzo (Enzo Life Sciences Inc., Farmingdale, NY) and horseradish peroxidase from Sigma. L-Fucose Assay Kit was purchased from Megazyme. All assay reagents were stored at −20 °C. The assays were carried out at room temperature.

### Bacterial strains and plasmids

*E. coli* NEB 5-alpha (New England Biolabs Inc., Ipswich, MA) was used for routine cloning and *E. coli* BL21 (DE3) was used for protein overexpression. Primers and synthetic DNA were purchased from Integrated DNA Technologies (IDT). Construction of plasmid pAfcA and pAfcB is reported in our previous work ^28^. Plasmids pAfcA and pAfcB comprises of a constitutive promoter and the fucosidase gene cloned onto a G9m-2 vector. Plasmid pET28:FDH was constructed by replacement of GFP in the pET28:GFP plasmid with the fucose dehydrogenase (FDH) gene with a His-tag fusion at the N-terminal. The FDH gene was amplified by PCR from the genomic DNA of *Bifidobacterium longum subsp. infantis* ATCC 15697™ (American Type Culture Collection, Manassas, VA). The FDH fragment was assembled into the pET28b vector, downstream of a T7 promoter, using NEBuilder® HiFi DNA Assembly Master Mix (New England Biolabs Inc., Ipswich, MA). The AlfA gene from *L. casei* was constructed by DNA synthesis (IDT) after codon-optimization for *E. coli* (IDT DNA Codon Optimization Tool). The gBlock was amplified and assembled into the G9m-2 vector, downstream of the constitutive promoter. Bacterial transformations were carried out using electroporation. The authenticity and orientation of the inserts was confirmed by DNA sequencing at the Iowa State DNA Facility (Ames, IA).

### Proof of concept using commercial kit

To test for the viability of using INT/PMS for a colorimetric readout for assaying free fucose, we used a L-Fucose Assay Kit (Megazyme Inc., Chicago, IL). Different concentrations of L-fucose were prepared and the assay carried out in a 96-well plate, according to the protocol supplied by the manufacturer. 2 μL of the dye mix was added to the reaction mixes and allowed to react for 5 mins. To test for the release of fucose from 2’-FL and 3-FL, a range of concentrations of the oligosaccharides were incubated with lysate from cells expressing pAfcA and pAfcB respectively, and followed by the kit protocol. The absorbance at 500 nm was monitored throughout the reaction.

### Enzyme isolation

The fucosidase and FDH expressing strains were cultured in 50 mL of LB medium at 37 °C at 250 rpm, overnight with appropriate antibiotics (carbenicillin, 50 ug/ml and 30 ug/ml kanamycin, respectively). The pET28:FDH was induced with IPTG (0.5 mM) at mid-log phase. These cultures were then incubated at 37 °C in a rotary shaker (250 rpm) for 16 hr. The overnight cultures were washed and resuspended in 2 mL water, and lysed by sonication (10 min, 30 s pulse, 30 s off, 30% amplitude). Debris was removed from lysates by centrifugation at 5000 xg for 30 mins and the supernatant stored at 4 °C. One unit of a-1,2-L-fucosidase activity was defined as the amount of enzyme required to release one mmol of L-fucose per minute from 2’-FL (2 mM) in sodium phosphate buffer (10 mM), pH 7 at 30 °C. The N-terminally His-tagged FDH was purified from the lysate under native conditions using the QIAexpress Ni-NTA Fast Start Kit (Qiagen, Vallencia, CA). 5 μl of each fraction was mixed with 5 μl of 2x Laemmli Sample Buffer (Bio-Rad, Hercules, CA). Each sample was heated at 95°C for 5 minutes and resolved on SDS-PAGE. The gel was visualized by Coomassie brilliant blue staining. The apparent molecular weight of FDH was approximately 32 kDa. The protein expression yields were calculated using a NanoDrop One Spectrophotometer (Thermo Scientific, Waltham, MA).

### Fabrication of paper-based sensor

The patterns for this study were printed on Whatman® grade 3 filter papers via wax-printing. They were purchased as sheets measuring 460 mm × 570 mm and cut to letter-size. Filter paper (Whatman #3) was chosen for its wicking properties and sturdiness, compared to filter papers of other thicknesses. The detection zone patterns were designed as circles with 7mm inner diameter and 1 mm thickness on Adobe Illustrator (Adobe, San Jose, CA). The detection zones were printed with a Xerox Color Qube 8570DN™ (Xerox, Norwalk, CT) printer using black hydrophobic wax-based ink.

The printed paper was placed in an oven set at 180 °C for 5 mins. To assess if the hydrophobic barriers fully penetrated the paper, water was spotted onto the detection zones to identify any bleeding and cross contamination between zones. If the water bled through, the paper was reheated and tested again till no bleed through was observed. The paper was used once cooled to room temperature. A layer of single-sided tape was used to seal the bottom side of the paper to contain the liquid within the barrier and prevent leakage.

### Performing paper-based assay

The lysates containing the enzymes were spotted at the center of the detection zones and allowed to dry. A 20 μL reaction system contained 2.5 uM NADP^+^, 1.25 mM INT (in DMSO) and 0.5 mM PMS (in DMSO). Table 1 shows a full listing of volumes of reagents required per reaction. A mastermix containing all the reagents excluding the enzymes and the dye was made and added to each well. The reactions were initiated by addition of the samples/standards, immediately followed by the addition of the dye reagent. Reactions were set up under low-light conditions. A color change was observed within 5 mins. All measurements were performed in triplicate. The paper was left to incubate in the dark till dry. Once dry, the paper was scanned using an Epson Perfection V800 Photo Scanner. To evaluate the limit of detection and generate a standard curve, a series of dilutions of 2’-FL/3-FL were made and analyzed. Data were fit via non-linear regression analysis using MATLAB.

**Table 1:** Procedure for running assay.

| Reagent | Standard | Blank | Sample |
| --- | --- | --- | --- |
| Distilled water | 10 μL | 11 μL | 10 μL |
| Buffer | 2 μL | 2 μL | 2 μL |
| NADP<sup>+</sup> | 0.5 μL | 0.5 μL | 0.5 μL |
| FDH | 0.1 μL | 0.1 μL | 0.1 μL |
| Fucosidase | 1 μL | 1 μL | 1 μL |
| **Sample/standard** | 5 µL | - | 5 µL |
| **INT/PMS dye** | 2 µL | 2 µL | 2 µL |

### Image and Statistical Analysis

The scanned images were analyzed on ImageJ that provided intensity data for each reaction. The MicroArray plugin was used that uses internally controlled regions of interest (ROI) and can measure microarray image stacks. The RGB channels were split and intensities were analyzed in the green channel and the mean values recorded. The background from the control with no analyte was calculated for each measurement. Calibration curves (**Figure 3C, 3D**) were built to get a quantitative readout from the measured transmittance. Comparison of our assay against a commercial microplate-reader was also carried out. We defined the LoD as the lowest fucose concentration that gave a colorimetric signal significantly above the control (no fucosylated analyte) sample (P ≤ 0.01). P values were determined by performing a paired parametric t test on Origin. To compare the analytical performance of our assay on biological fluids, we analyzed 6 random trials, with one half of the experimental data set allocated toward blind testing.

### Long term stability of the paper-based assay

To assess the stability of the paper assay, the AfcA enzyme lysates were spotted onto the detection zones and left to air-dry in the fume hood for an hour at room temperature. Another batch of spotted paper was subjected to lyophilization overnight. The papers were either stored at room temperature or at 4 °C in plastic bags with silica pouches in the dark. To determine the enzymatic activity after specific storage time, the assays for detection of 2’-FL were carried out over a period of 45 days by rehydration with standard solutions. The colorimetric output was analyzed as described earlier. All measurements were carried out at room temperature and in triplicates.

### Analysis of biological samples

Pooled, deidentified human breast milk samples were purchased from BioIVT (Westbury, NY). Surine™ Negative Urine Control was purchased from Cerilliant (Cerilliant Co., Round Rock, TX). HMOs were extracted from human breast milk using established procedures. 200 μL of milk was centrifuged at 9,000 rpm for 20 mins at 4 °C to remove lipids. After the top layer was removed, 400 μL of ethanol was added and centrifuged at 9,000 rpm for 10 mins at 4 °C and the supernatant collected. The ethanol was removed by vacuum centrifuging and reconstituted in water to different concentrations before analysis using the assay and validation of relative abundance carried out using LC-MS/MS. We used an Agilent Technologies 1100 Series HPLC system coupled to an Agilent Technologies Mass Selective Trap SL detector (Agilent Technologies, Santa Clara, CA) equipped with an electrospray ion source (ESI) and controlled by ChemStation A.10.02, including data acquisition and processing. Both MS and MS/MS spectra were acquired in the negative-ion mode with an acquisition rate of 1 ◻s per spectrum over ranges of m/z 300 to 2,000 (for MS) and m/z 50 to 2,000 (for MS/MS). Precursor-ion selection was performed automatically by the data system based on ion abundance. Three precursors were selected from each MS spectrum to carry out product-ion scanning. Products were analyzed on a Prevail Carbohydrate ES (250 mm × 4.6 mm id; 5 μm; Alltech Associates, Deerfield, IL). The mobile phase consisted of water (aqueous solution containing 5 mM ammonium acetate) and acetonitrile (AcN containing 5 mM ammonium acetate). The solvent gradient was delivered at a flow rate of 1 ◻ml/min and consisted of an initial linear increase from 85% to 50% AcN over 10 min, held at 50% AcN for 2 mins and followed by a linear decrease to starting conditions for 2 min. The injection volume was 5 ◻μl. The dynamic range and linearity for the quantification of 2’-FL and 3-FL were evaluated through the use of a series of dilutions as standard curves. To analyze 3-FL, the samples were concentrated in a vacuum centrifuge (Eppendorf, Hauppauge, NY). Disparities with other cited literature may be due to different analytical technique and differences in biochemical extraction process.

