## Supplemental Information for "Paper-based Sensing of Fucosylated Biological Compounds"

**Supplementary Information**

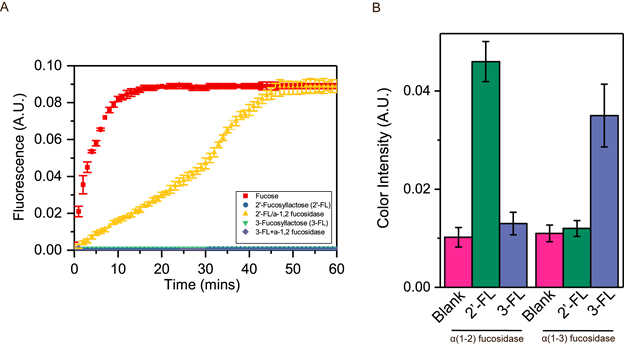

**Figure S1: Orthogonality of enzymatic assay. (A)** Timecourse of 340 nm absorbance as measured on a micro-plate reader with fucose (red), 2’-FL (blue) and 3-FL (green) in the absence of α-1,2-fucosidase and 2’-FL (yellow) and 3-FL (purple) in the presence of α-1,2-fucosidase. **(B)** Test for orthogonality of fucosidases based on measurement of colorimetric readout of assay.

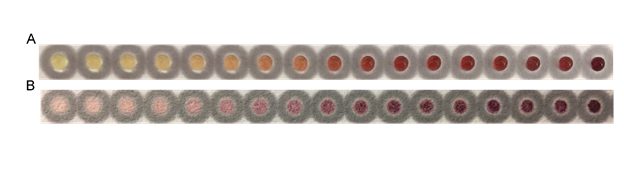

**Figure S2: Colorimetric readout using paper-based assay. (A)** Image showing dilutions of commercial 2’-FL captured within seconds of reaction. **(B)** Image after the reaction were dry.

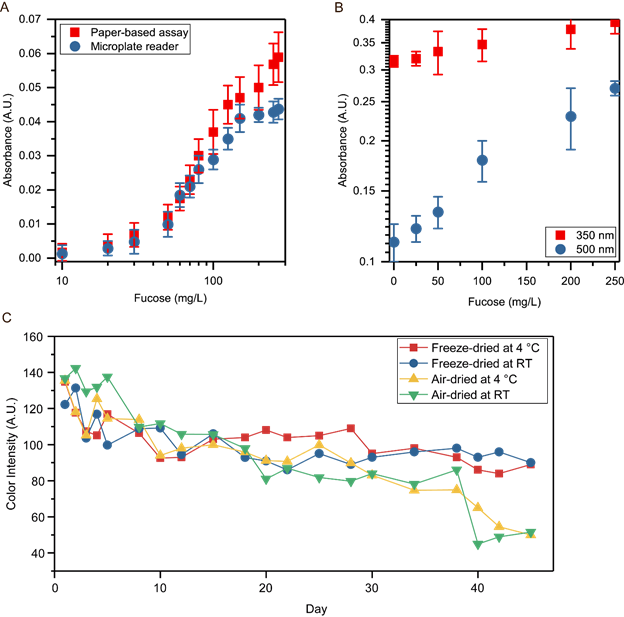

**Figure S3: Testing the robustness of the assay and paper device. (A)** Colorimetric readout using imaging software in comparison to readouts using a microplate reader. Based on the Mann–Whitney–Wilcoxon test, the two datasets in the range 0-75 mg/L, are not significantly different. **(B)** Sensitivity of tetrazolium dye (500 nm) in comparison to NADPH absorbance (350 nm). Mean readings with error bars representing standard deviations of three independent experiments is shown. **(C)** Viability of paper device after 45 days under different enzyme immobilization techniques and storage conditions.

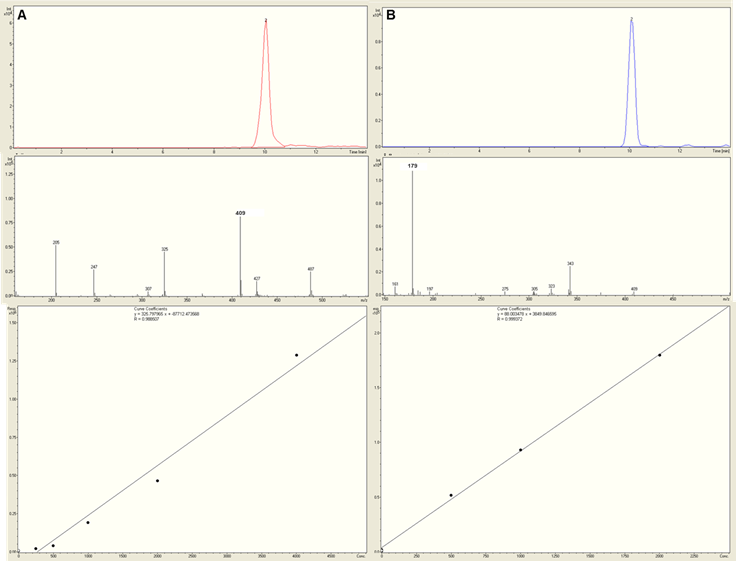

**Figure S4: Quantification of fucosyllactose isomers using HPLC-MS. (A)** *(top to bottom)* Extracted ion chromatograms of 2’-FL (m/z 523); specific ion profile for 2’-FL from MS; a standard curve was created by LC-MS/MS analysis with varying levels of commercial 2’-FL. **(B)** *(top to bottom)* Extracted ion chromatograms of 3-FL (m/z 523); specific ion profile for 3-FL from MS; a standard curve was created by LC-MS/MS analysis with varying levels of commercial 3-FL.

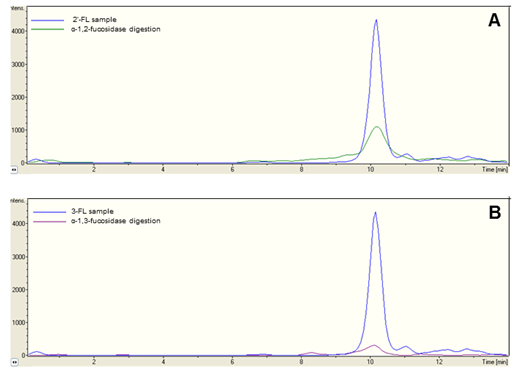

**Figure S5: Validation of isomer peaks by exofucosidase digestion. (A)** Peaks before and after treatment with α-1,2-fucosidase to digest 2’-FL **(B)** Peaks before and after treatment with α-1,3-fucosidase to digest 3-FL.

**Abbreviations**

| ADCC | Antibody-Dependent Cellular Cytotoxicity |
| --- | --- |
| CE | Capillary electrophoresis |
| 2’-FL | 2’-fucosyllactose |
| FDH | Fucose dehydrogenase |
| HMO | Human milk oligosaccharides |
| HPAEC-PAD | High-Performance Anion-Exchange Chromatography with Pulsed Amperometric Detection |
| HRP | Horseradish peroxidase |
| IgG | Immunoglobulin G |
| INT | 2-(4-iodophenyl)-3-(4-nitrophenyl)-5-phenyl-2H- tetrazolium chloride |
| LPS | Lipopolysaccharide |
| NADP | Nicotinamide adenine dinucleotide phosphate |
| PDMS | Polydimethylsiloxane |
| 4-PL | 4-parameter logistic model |
| PLA2 | Phospholipase A2 |
| PMS | Phenazine methosulfate |
| PNGase F | Peptide-N4-(N-acetyl-beta-glucosaminyl) asparagine-amidase |
